# VESNA: An Open-Source Tool for Automated 3D Vessel Segmentation and Network Analysis

**DOI:** 10.1101/2025.03.05.641600

**Authors:** Magdalena Schüttler, Leyla Doğan, Jana Kirchner, Süleyman Ergün, Philipp Wörsdörfer, Sabine C. Fischer

## Abstract

**Background:** Vasculature is an essential part of all tissues and organs and is involved in a wide range of different diseases. However, available software for blood vessel image analysis is often limited: Some only process two-dimensional data, others lack batch processing, putting a time burden on the user, while still others require tightly defined culturing methods and experimental conditions. This highlights the need for a software that has the ability to batch process three-dimensional image data and requires few and simple experimental preparation steps.

**Results:** We present VESNA, a Fiji (ImageJ) macro for automated segmentation and skeletonization of three-dimensional fluorescence images, enabling quantitative vascular network analysis. It requires only basic experimental preparation, making it highly adaptable to a wide range of possible applications across experimental goals and different tissue culturing methods. The macro’s potential is demonstrated on a range of different image data sets, from organoids with varying sizes, network complexities, and growth conditions to expanding to other 3D tissue culturing methods with an example of hydrogel-based cultures.

**Conclusions:** With its ability to process large amounts of 3D image data and its flexibility across experimental conditions, VESNA fulfills previously unmet needs in image processing of vascular structures and can be a valuable tool for a variety of experimental setups around three-dimensional vasculature, such as drug screening, research in tissue development and disease mechanisms.

## 1. Introduction

In recent years, three-dimensional tissue culture techniques, such as organoids, have emerged as powerful tools for studying organ development, function, and diseases. Blood vessel organoids are developed by selforganization like vasculogenesis. Their organization and branching pattern mirrors the tissue vascularization during the embryogenic development. Beyond its significance in cardiovascular diseases, the vascular system is central to conditions like diabetes [1] and tumor progression [2, 3]. Recent efforts have extended to exploring assembloids and vascularized organoids, enabling the creation of older, larger, and more complex models [3, 4, 5, 6].

While significant numbers of organoids can be produced and imaged with relative ease, the manual image processing of the resulting data is often a limiting factor [7, 8].

Early open-source plugins and software tools, such as *AngioQuant* [9], *VESGEN 2D* [10], and *AngioTool* [11] provide semi-automated and automated analyses of two-dimensional images. Over time, newer tools like *Quantification* [12], *REAVER* [13], *Angiogenesis Analyser* [14], and *Q-VAT* [15] have introduced advanced functionalities. *SproutAngio* [16] and *VesselExpress* [17] further include the ability to process three-dimensional images, a feature that is essential for organoid analyses.

Nevertheless, these software tools exhibit notable limitations when applied to blood vessel organoids. For example, *SproutAngio* lacks the capacity for batch processing, making image processing and analyses time-consuming, while *VesselExpress* has been optimized for hydrogel-filled vessels, which is challenging to achieve in organoids. Specialized tools such as *Automated Sprout Analysis* often demand narrowly defined experimental setups and additional staining protocols such as F-actin and nucleus stains, which may not align with the experimental goal.

These limitations highlight a clear gap, which we address with our macro VESNA (Vessel Segmentation and Network Analysis). Designed explicitly for three-dimensional fluorescence images of vascular networks, VESNA requires only a vasculature-specific stain, such as CD31. To ensure cross-platform functionality and ease of use, it is built for the open-source platform Fiji (ImageJ) [18]. Our macro emphasizes batch processing and minimal user intervention, enhancing efficiency and reducing processing time.

We validated VESNA’s performance against ground truth data and evaluated its sensitivity to parameter settings. Furthermore, we demonstrated its ability to process diverse datasets, including drug-treated organoids, organoids cultured under varying growth conditions, and hydrogel-based models. Regardless of image size, network complexity, or organoid age, the macro effectively analyzed vascular network size and structure. This versatility underscores its value in diverse experimental setups, addressing a critical need in vascular organoid research.

## 2. Materials and Methods

### 2.1 Image Processing and Data Analysis

The development of our macro VESNA (Vessel Segmentation and Network Analysis) builds upon the foundational work of *Quantification* by Rust *et al*. [12]. *Quantification* is a macro for the image processing software Fiji (ImageJ) [18], that allows for automated segmentation of two-dimensional fluorescence microscopy images of vascular networks. VESNA expands on this by enabling the processing of three-dimensional image data that better reflect the complexity of vascular structures *in vitro*, thereby ensuring a more comprehensive representation of biological systems and processes. We included a number of pre- and post-processing steps to yield high quality image and measurement data, and introduced batch processing to further automatize the image processing workflow, minimizing user involvement during the processing and improving efficiency.

The macro’s image processing workflow can generally be divided into three steps: First, the raw image is preprocessed to ensure the fidelity of later steps. The image is then binarized to define the areas of the vascular structures. Finally, skeletonization simplifies the binary image for measurement (Figure 1).

**Figure 1.**
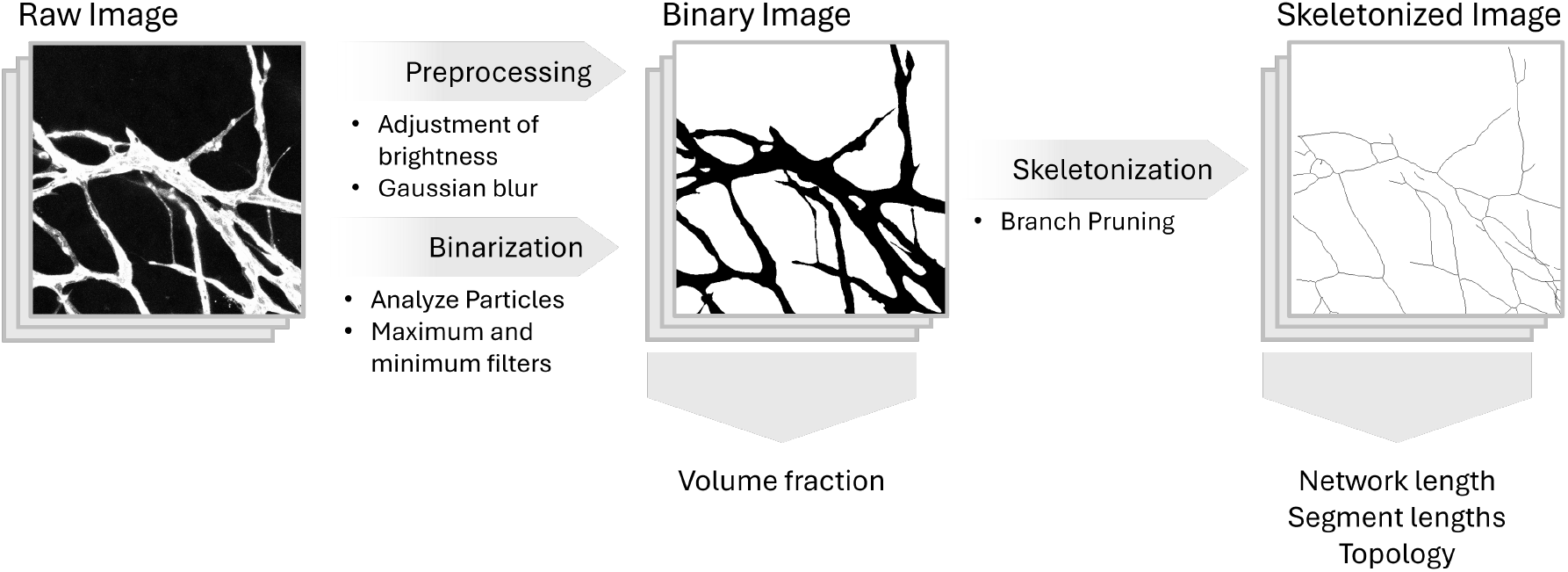
Schematic overview of the three main steps of VESNA’s image processing workflow, the associated postprocessing methods, and the measurements carried out by the macro. Depicted are cropped Z maximum projections of an image of Dataset D.

#### Preprocessing

During preprocessing, several operations are applied to the raw image to reduce noise and achieve high quality of the following binarization. First, the brightness of the image is adjusted to ensure the detection of vessels with low fluorescence intensity. To compensate for the highly heterogeneous fluorescence within the available images, a three-dimensional Gaussian blur is utilized. This ensures the recognition of the vessels as a whole instead of a falsely fragmented segmentation. Both steps can be adjusted by user-defined parameters, using the interface of the macro. The default values of these and all other parameters are listed in Table S1.

#### Binarization

Once the preprocessing steps are completed, the image is binarized to identify the areas of the vessel structures. For this, the Yen threshold [19] is used. To improve the fidelity of the resulting binary image, the macro first removes artifacts below a defined pixel parameter value using the Analyze Particles function before applying three-dimensional maximum and minimum filters to connect fragmented vessel structures. Lastly, internal holes in the binary image are removed by the Fill Holes function included in the package *MorphoLibJ* [20]. This step is essential to prevent artifacts in the skeleton that would otherwise be hard to remove (see Figure 2 for an example).

**Figure 2.**
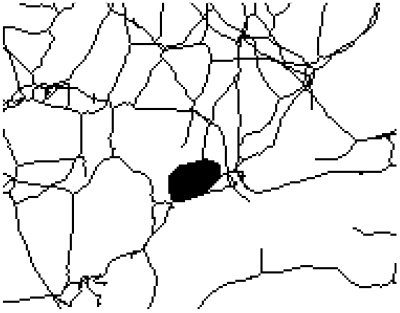
Exemplary cropped Z maximum projection of a skeleton to display the artifacts that are removed by the Fill Holes function. The skeleton was generated from Dataset A.

The resulting binary image is used to calculate the volume fraction of the acquired capillary network. All other measurements are performed based on the skeletonized image.

A machine learning-based binarization via *Trainable Weka Segmentation* [21] was tested as an alternative to the binarization using preinstalled thresholding methods, but was found to require too much processing power at original image dimensions. Scaling of the images led to significantly worse segmentation results. For this reason, less intensive automated binarization by Yen thresholding was selected to ensure high quality results while limiting the required processing power and time.

#### Skeletonization

The binary image is skeletonized to simplify the vessel structures for network analyses. This produces a network of one voxel thick segments connected to each other by junction points and ended by end points. For this, the plugin *Skeletonize3D* [22] is utilized. Small artifactual network segments are removed by a version of the BeanShell script *Prune Skeleton Ends* [23], that we made compatible with our macro while maintaining its functionality.

Finally, the remaining measurements are performed. The length of the entire vessel network is measured in voxels and voxels per mm^3^ using the plugin *Analyze Regions 3D*, that is part of the *MorphoLibJ* package [20]. The plugin *AnalyzeSkeleton* [24] allows access to several measurements, including the number of skeletons, segments, junction and end points, as well as the segment length.

In this context, a segment is defined as a part of a skeleton that ends in two junction points, two end points, or one junction point and one end point (Figure 3). A skeleton is a group of segments that are directly connected to each other. Under this definition, the skeletonized image of one vascular network, for example of one blood vessel organoid, could—and often does—consist of several skeletons. However, the term skeleton is often used inconsistently for the entire skeletonized image. In order to avoid confusion, we will only use the term skeleton to refer to the former definition.

**Figure 3.**
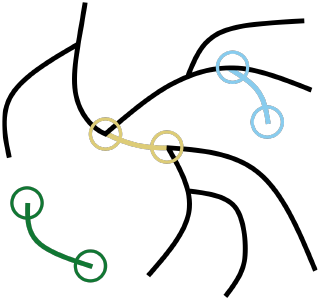
Illustration for explanation of the terms skeleton and segment. Exemplary segments are highlighted in colors, circles mark junctions and end points. A skeleton is a group of segments that are directly connected to each other.

#### Parameter Optimization

For all data sets in this work, we established an optimized parameter setting that was based on manual comparison between the raw, the binary and the skeletonized image.

Proper setting of the brightness parameters were verified by comparison of the binary image with the raw image. Ideally, the entire vessel structure should be recognized during the segmentation, without loss of the definition in small and detailed structures. Recognition of weakly fluorescing vessels can be improved by lowering the brightness maximum. Conversely, to improve definition, the brightness minimum and maximum can be increased. In practice, particularly in images exhibiting non-homogeneous fluorescence and higher background fluorescence, achieving a balance is often challenging. In such cases, implication of the Subtract Background function can lead to better results. The definition of the vessels with higher fluorescence should usually be the priority. While this may lead to weakly fluorescing vessels not being recognized in their entirety, this approach can still yield accurate network measurement data for the recognizable regions. In any case, the priority was set according to the specific goals of the experiment at hand.

The main function of the maximum and minimum filters, as well as the Gaussian blur, is to recombine fragmented vessels in the skeletonized image. Additionally, the Gaussian blur removes some small artifactual segments, that occur at places with irregularly shaped vessel structures in the binary image (Figure 4). Therefore the setting of these filters was assessed by comparing the skeleton to the raw image. The parameter values were chosen so that such artifacts or fragmented vessels were minimal, while the detail of the network was maintained.

**Figure 4.**
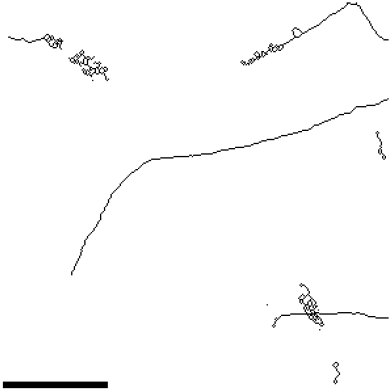
Exemplary image showing artifacts resulting from incorrectly set *σ* parameter for Gaussian blur. Depicted is a cropped Z maximum projection of a skeletonized image of Dataset D that was generated with the omission of the Gaussian blur. Scale 50 µm.

Verification of the pixel threshold value for the Analyze Particles function was achieved by checking for small artifacts in the binary or skeletonized image that are not directly connected to larger structures.

Branch Pruning was used to remove any remaining short artifactual segments that are directly connected to the network structure. The length threshold was considered correctly if the skeleton did not show any such short artifacts.

### 2.2 Generation of Sample Data

To assess the functionality of VESNA on different types of data, vascular networks were generated in blood vessel organoids and hydrogel-based tissue cultures of varying cell count and age. Additionally, treatment with antiangiogenic substances was performed to demonstrate the possible usage of VESNA in drug screening assays.

#### Human iPSC Culture

Human induced pluripotent stem cells (hiPSCs) were generated from commercially available normal human dermal fibroblasts (Promocell, Heidelberg, Germany) by reprogramming using the hSTEMCCA-lentiviral construct [25, 26]. The hiPSCs are cultured on human embryonic stem cell (hESC)-qualified Matrigel (Corning, New York, NY, USA)-coated culture plates in StemMACS iPS Brew medium (Miltenyi Biotec, Bergisch Gladbach, Germany). The culturing medium is replaced daily. For passaging, cells are dissociated at 80 % confluency with StemPro Accutase (Thermo Fisher Scientific, Waltham, MA, USA) for 5 min at 37 °C to obtain a single-cell suspension. Subsequently, hiPSCs are replated in StemMACS medium, supplemented with 10 nM thiazovivin (TZ) (LC Labs, Woburn, USA).

#### Blood Vessel Organoid Generation

Blood vessel organoids are self-organizing 3D tissue cultures derived from hiPSCs. They consist of a loose mesenchymal connective tissue harboring a hierarchically organized and branching endothelial network. Different approaches were used to generate blood vessel organoids for this study. For Dataset A, blood vessel organoids were generated following a previously established protocol [27]. In brief, hiPSCs are detached from the cell culture dish using StemPro Accutase for 5 min at 37 °C to obtain a single-cell suspension. Subsequently, 4000 cells per 100 µL StemMACS iPS Brew medium (Miltenyi Biotec, Bergisch Gladbach, Germany) supplemented with 10 mM TZ are prepared and pipetted into each well of an agarose-coated 96-well plate. Cells are cultured for 24 h at 37 °C and 5 % CO_2_ in a humidified incubator to allow hiPSC aggregate formation. After 24 h, the medium is changed to mesodermal induction medium (MIM) (Advanced DMEM/F12 (Gibco/Life Technologies, Westham, MO, USA) 100 %, L-Glutamine (Gibco/Life Technologies, Westham, MO, USA) 0.2 mM, Ascorbic acid (Sigma-Aldrich, St. Louis, MO, USA) 60 µgµL^*−*1^, CHIR 99021 (Sigma-Aldrich, St. Louis, MO, USA) 10 µM, BMP4 (PeproTech, Cranbury, NJ, USA) 25 ngmL^*−*1^) for 72 h. After that, the medium is discarded, and organoids are cultivated in vascular growth medium (VGM) (Neurobasal medium/DMEM-F12 (Gibco/Life Technologies, Westham, MO, USA) (50 %/50 %), B27 without Vitamin A (Gibco Life Technologies, Westham, MO, USA) 1×, N2-Supplement (Gibco Life Technologies, Westham, MO, USA) 1×, L-Glutamine (Gibco Life Technologies, Westham, MO, USA) 2 mM, Ascorbic acid 60 µgµL^*−*1^, vascular endothelial growth factor (VEGF) (ProteinTech, Rosemont, IL, USA) 100 µgµL^*−*1^). After 48 h, the medium is changed to organoid maintenance medium (OMM) (Neurobasal medium/DMEM-F12 (50 %/50 %), B27 without Vitamin A 1×, N2-Supplement 1×, L-Glutamine 2 mM, Ascorbic acid 60 µgµL^*−*1^) and organoids are cultured at 37 °C and 5 % CO_2_ in a humidified incubator on a cell culture rocking table. The medium is changed every other day. Vascular networks were analyzed at day 14.

For Datasets B and C, the same protocol was used with the exception that hiPSCs are seeded into custom-made aggrewell-like agarose micromolds instead of agarose-coated 96-wells. To prepare agarose micromolds, 2 % agarose (Biozym, Hessisch Oldendorf, Germany) was boiled in water and hot agarose solution was poured into the well of a 12-well culture dish containing a 3D-printed negative master mold. Two different types of negative master molds were used (either with 159 micropillars to create small agarose molds (159 × 700 µm diameter) or with 31 micropillars to create larger agarose molds (31 × 2 mm diameter)). After one hour, the negative master mold was removed from the solidified agarose and the obtained micromold was placed into a well of a 12-well plate (small molds) or 6-well plate (large molds). 2-10 *×*10^4^ cells are seeded per small micromold and 1-2 *×*10^5^ cells for large molds and cultured for 24 h in StemMACS iPS Brew medium (Miltenyi Biotec, Bergisch Gladbach, Germany) at 37 °C and 5 % CO_2_ in a humidified incubator. 159 iPSC aggregates form in small molds and 31 iPSC aggregates in large molds. After 24 h, the medium was changed to MIM (Advanced DMEM-F12 (Gibco/Life Technologies, Westham, MO, USA) 100 %, L-Glutamine (Gibco/Life Technologies, Westham, MO, USA) 2 mM, Ascorbic acid (Sigma-Aldrich, St. Louis, MO, USA) 60 µgµL^*−*1^, CHIR 99021 (Sigma-Aldrich, St. Louis, MO, USA) 10 µM, BMP4 (Pe-pro Tech, Cranbury, NJ, USA) 25 ngmL^*−*1^) for 72 h. After that, the organoids were removed from the micromolds and transferred into a well of a 6-well plate. The organoids were then cultured in VGM (Neurobasal medium/DMEM-F12 (Gibco/Life Technologies, Westham, MO, USA) (50 %/50 %), 5 % heat-treated fetal calf serum (FCS), B27 without Vitamin A (Gibco Life Technologies, Westham, MO, USA) 1×, N2-Supplement (Gibco Life Technologies, Westham, MO, USA) 1×, L-Glutamine (Gibco Life Technologies, Westham, MO, USA) 2 mM, Ascorbic acid 60 µgµL^*−*1^, VEGF (ProteinTech, Rosemont, IL, USA) 50 µgµL^*−*1^) on a cell-culture rocker to avoid aggregation.

For antiangiogenic drug treatment, organoids from small molds were separated into two comparable groups at culture day 8. Subsequently, one group is treated for 72 h with 15 nM sorafenib (Gibco, Karlsruhe, Germany). For the control group, sorafenib is omitted. Analyses of vascular networks were performed after 72 h of sorafenib treatment (culture day 11).

For Dataset B, organoids from small and large molds were compared at culture day 8.

For visualization of vessel networks, whole-mount immunofluorescence staining and ethyl cinnamate-based tissue clearing were performed as previously described [28]. A primary antibody directed against CD31 (PECAM1) (Agilent, Santa Clara, CA, USA, mouse, M0823, 1:200) was used to detect endothelial cells. Cy3-conjugated secondary antibodies (Dianova, Hamburg, Germany) were used to visualize the primary antibody. Imaging was performed using the Nikon Eclipse Ti confocal laser scanning microscope (Nikon, Tokyo, Japan) with a long working distance air objective (20×) for taking z-stack images. Nikon NIS Elements Confocal software version 4.13.05 (Nikon, Tokyo, Japan) was used for imaging.

#### Generation of Bioprinted Hydrogel-Based Vascular Cultures

HiPSCs-derived mesodermal progenitor cells (MPCs) can spontaneously form a loose mesenchymal connective tissue harboring a branching endothelial network after extrusion-based bioprinting using Matrigel as a bioink, as previously reported [29].

For Dataset D, hiPSCs were first converted to MPCs using an adherent 2D cell culture protocol [30]. Therefore, 3 *×*10^5^ hiPSCs*/*cm^2^ were seeded per Matrigel-coated well of a 6-well plate and cultured for 24 h in StemMACS iPS Brew medium supplemented with 10 mM TZ. Afterwards, the cells were cultured for 3 days in MIM (Advanced DMEM/F12 100 %, L-Glutamine 0.2 mM, Ascorbic acid 60 µgµL^*−*1^, CHIR 99021 10 mM, BMP4 25 ngmL^*−*1^) at 37 °C and 5 % CO_2_.

At day 4, cells were dissociated using Accutase. 6 *×*10^6^ cells were mixed with cold Matrigel solution (3.6 mgmL ^*−*1^ final protein content in Advanced DMEM/F12) and a homogeneous gel-cell mixture (bioink) was prepared. The mixture was extruded into silicon molds through a 22G _1_/_4_ inch lock tip nozzle under a continuous pressure of 10 kPa, using the VIEWEG GmbH brand DC 200 model analog dispenser to form gel disks with a volume of 100 µL. The printed disks were kept at room temperature for 45 min. Subsequently, cell-loaded gel disks were washed with PBS and transferred into a cell culture flask containing vascular differentiation medium (Advanced DMEM/F12 100 %, 5 % heat-treated FCS, 1 % Penicillin/Streptomycin (P/S), TZ 10 mM, L-Glutamine 0.2 mM, Ascorbic acid 60 µgµL^*−*1^, hVEGF-A 60 µgµL^*−*1^). For whole-mount immunofluorescence analyses, the disks were removed from the culture at days 7 and 17 and washed three times with PBS for 5 min. The disks were fixed with 4 % paraformaldehyde (PFA) in PBS for 24 h at 4 °C and washed again three times with PBS for 30 min to remove residual PFA.

After washing, the disks are incubated in blocking buffer (PBS, 4 % normal goat serum (NGS, G9023, Sigma-Aldrich, USA), 0.1 % Triton X-100) for 3 h at 4 °C. The samples are incubated with a primary antibody directed against CD31 (PECAM1) overnight at 4 °C. Afterwards, samples were rinsed three times with PBS and exposed to Cy3-conjugated secondary antibodies to visualize the primary antibody (Dianova, Hamburg, Germany). Imaging was performed using the Nikon Eclipse Ti confocal laser scanning microscope (Nikon, Tokyo, Japan) with a long working distance air objective (20×) for taking z-stack images. Nikon NIS Elements Confocal software version 4.13.05 (Nikon, Tokyo, Japan) was used for imaging.

### 2.3 Statistical Analysis

For hypothesis testing, we used the two-sided Mann-Whitney U test with a significance level of *α∗* = 0.05 and Bonferroni correction. The Bonferroni-corrected significance levels *α* are listed in Table 1. In the case of the Sensitivity of Parameters (Section 3.2), the Branch Pruning parameter has no effect on the volume fraction and is thus omitted from the test in this measurement, leading to a separate *α* value.

**Table 1.** Bonferroni-corrected significance levels *α* calculated for all statistical tests performed.

| section | measurements | $\alpha$ |
| --- | --- | --- |
| Sensitivity of Parameters<br>(Section 3.2) | volume fraction | 0.0029 |
|  | other | 0.0036 |
| Application to Sample Data<br>(Section 3.3) | all | 0.050 |

Additionally, the corresponding effect sizes are tested using the Glass’s Δ. It is calculated with 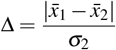, where 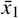 and 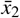 are the mean values of the compared groups 1 and 2, and *σ*_2_ is the standard deviation of the untreated or control group 2. The resulting Glass’s Δ values give insight into the extent of the deviation. Reflecting Cohen [31], a value of Δ *≤* 0.2 indicates a low effect size, 0.2 *<* Δ *≤* 0.5 a medium effect size, and Δ *>* 0.5 a high effect size. The statistical analyses were implemented in Python Version 3.11.7.

### 2.4 Data availability

The macro together with instructions and a sample image are available on https://github.com/scfischer/schuettler-et-al-2025/. Upon publication, we will also provide it on Zenodo.

## 3. Results

### 3.1 Image segmentation quality

We evaluated the segmentation quality of VESNA using an example image. Specifically, we quantified the overlap between the automatically segmented image and a manually drawn ground truth. For this analysis, we processed a section of an image of a blood vessel organoid (Section 2.2, Dataset A), using the parameter settings in Table S1. The ground truth was created by manually drawing an ideal skeleton into each slice of the raw image stack (Figure 5). We then compared the segmentation results with the ground truth and with a dilated ground truth to include a less stringent quantification. Despite the apparent increase in area, the dilation preserved most of the detail and definition of the ground truth (Figure 6).

**Figure 5.**
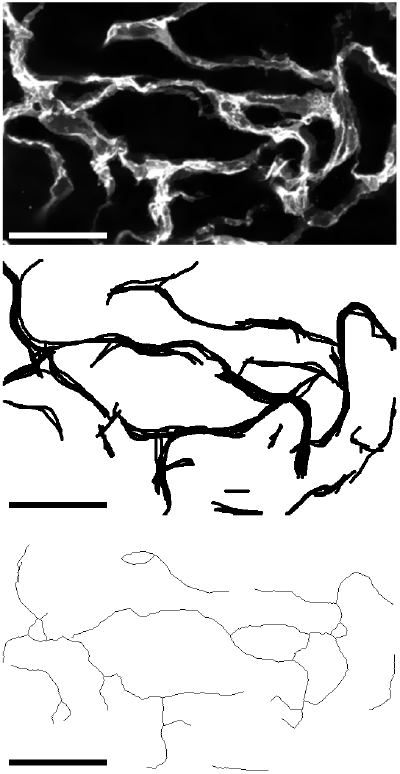
Exemplary cropped Z maximum projections of (from top to bottom) the fluorescence microscopy image of a blood vessel organoid, the ground truth, and the skeletonized image generated by VESNA based on Dataset A. Scale 50 µm.

**Figure 6.**
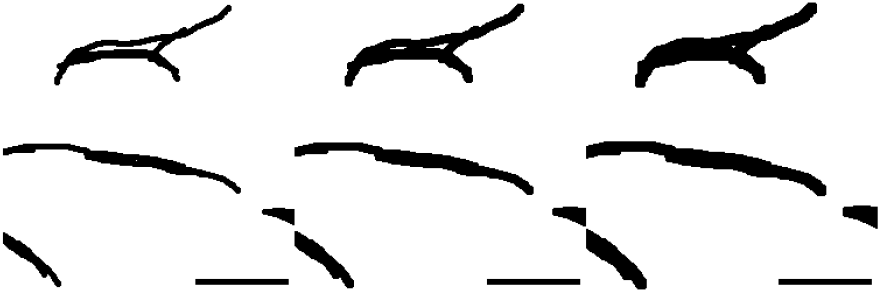
An exemplary cropped Z maximum projection of the ground truth of Dataset A: (from left to right) original, dilated once, and dilated twice. Scale 25 µm.

The resulting data indicate a noticeable overlap between the skeletonized image and the ground truth (Table 2), increasing with the level of dilation applied. A visual comparison of the ground truth and the skeletonized image provides insights into potential factors influencing this outcome.

**Table 2.**
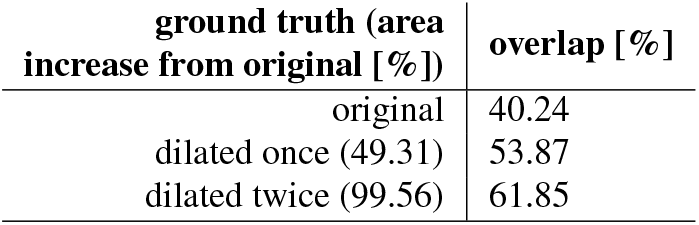
Percentages of skeleton voxels overlapping with the original and dilated ground truth, as well as area increase of the dilated ground truth from the original.

The skeleton is a one-voxel-wide structure, making quantitative comparison with the ground truth highly sensitive. Discrepancies may arise due to slight displacements of recognized vessel structures between VESNA’s output and the ground truth, as well as inaccuracies in the ground truth itself caused by drawing errors or ambiguous structures in the original image (Figure 7).

**Figure 7.**
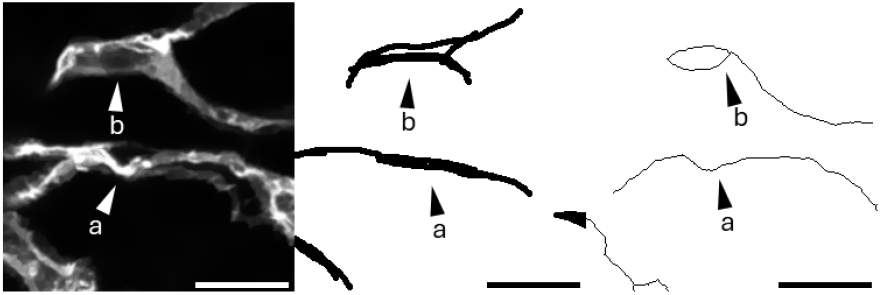
Exemplary cropped Z maximum projection of (from left to right) the raw image of a blood vessel organoid, the ground truth, and the skeletonized image of Dataset A. Marked is an example of a region where the macro recognized a structure that is dislocated relative to the ground truth (**a**), and one where the segmented structures are more detailed than defined by the ground truth (**b**). Scale 25 µm.

We further recognised low-fluorescence structures in the original image. Many of these structures can be identified or completed by eye with relative ease, depending on the fluorescence intensity, but are sometimes not detected by VESNA. The adjustment of the brightness parameters as a way to recognize these structures leads to a loss of detail in finer and brighter fluorescence structures in this image. This implies a trade-off that is discussed further in Section 2.1.

We conclude that even though the degree of overlap of VESNA’s segmentation with the ground truth appears moderate at first glance, a closer comparison of the skeletonized image, the ground truth and the raw image indicates a satisfactory result.

Next, we investigated the sensitivity of the overlap between the skeletonized image and the ground truth to the parameter settings of VESNA (Table S2). As a basis, we used parameter values that were optimized manually as detailed in Section 2.1.

The resulting percentages of overlapping skeleton voxels relative to the total skeleton voxels show that the deviations of the overlaps are small (Figure 8). The most prominent decrease in overlap is caused by lowering the brightness maximum parameter. This setting leads to increased false positive segmentation, causing a higher number of skeleton voxels (7108 voxels compared to 4674 voxels resulting from optimized settings) and leading to the decreased percental overlap (32.96 % with the original ground truth compared to 40.24 %). The largest increase of the overlap is produced by a higher *σ* value for the Gaussian blur. However, the effect is minimal (42.72 % overlap with the original ground truth compared to 40.24 % resulting from optimized settings).

**Figure 8.**
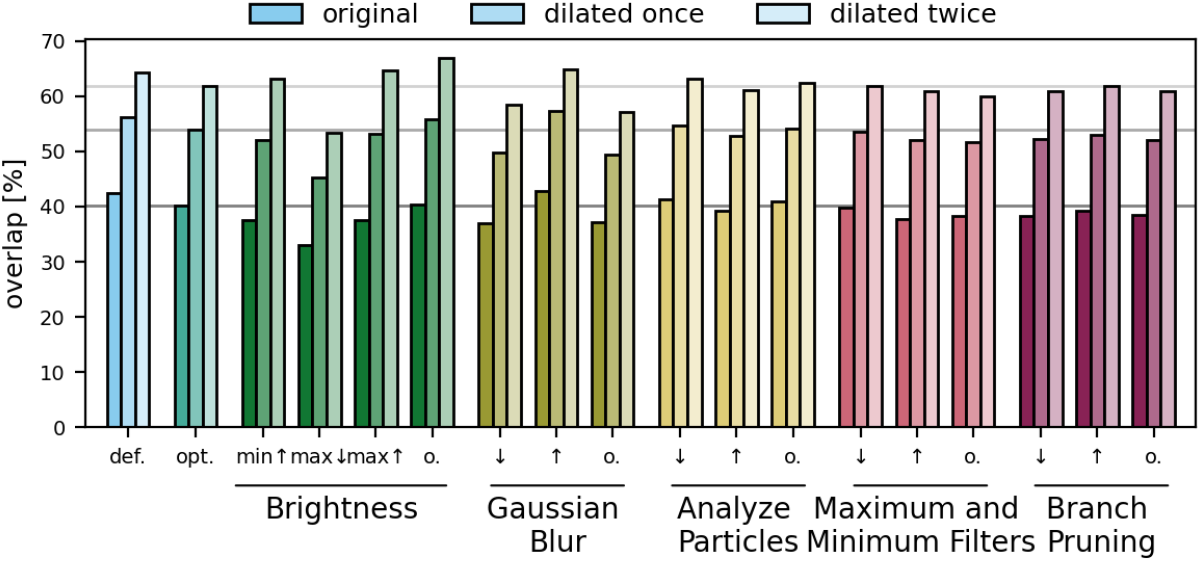
Percentages of skeleton voxels of Dataset A overlapping with the original and dilated ground truth at varying parameter settings (sample size *n* = 1). Horizontal lines mark the overlap values with optimized parameter settings for easier comparison.

The results demonstrate the robustness of the overlap between the resulting skeletonized image and the ground truth across varying parameter settings for this dataset.

Our results highlight the challenges of quantitatively evaluating segmented images generated by VESNA. Therefore, this process is not repeated for other raw images. Instead, we emphasize the significance of visually assessing the processed image data. In addition, we investigate the effect of the parameter settings on the network measurements.

### 3.2 Sensitivity of Network Measurements

To determine the impact of parameter adjustments on the network measurement data, we performed a sensitivity analysis. A subset of the sample data that was found to be computationally viable for the high number of processing runs necessary for this analysis was selected (Section 2.2, Datasets C and D). The individual functions of the script are tested with parameter values chosen to be just outside the range that is typically found to be useful for most image data, as well as with the function omitted (Table S2). Notably, the actually useful range may vary.

The deviations between the network measurements generated from the tested parameter settings and the corresponding optimized settings are mostly not statistically significant (Figure S1). A notable exception is the segment length, which shows a significant deviation in 22 out of 34 parameter setting groups. This is easily explained by the large amount of data points generated in this measurement. Given that each image contains hundreds of segments, which are measured and plotted individually, a statistical significance is much more attainable.

Although significant deviations in the other measurements could not be detected, the effect sizes (Table S3) indicate several larger differences. These are generally expected and are explained by considering each function’s purpose. For instance, a low *σ* value or the omission of the Gaussian blur frequently results in failed or incomplete recombination of fragmented vessel structures. This in turn leads to an increased number of segments and skeletons, as seen in Figure 9 and Table 3. Conversely, a high *σ* value reconnects many of the fragmented vessel structures but leads to a loss of detail indicated by a decrease in segment and skeleton numbers.

**Figure 9.**
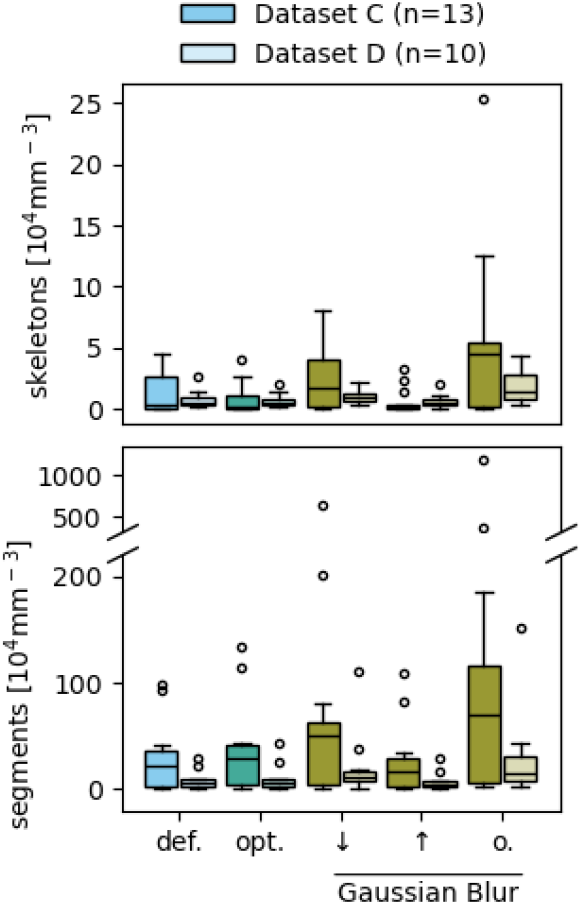
Comparison between the measurement data resulting from default (**def**.) and optimized (**opt**.) parameter settings, as well as from varying *σ* values (***↓, ↑***) and the omission (**o**.) of the Gaussian blur of Datasets C and D. None of the depicted data show a statistically significant deviation below a Bonferroni-corrected significance level of *α* = 0.0036 from the optimized data. Complete data are shown in Figure S1.

**Table 3.** Effect sizes of the deviation of the numbers of skeletons and segments resulting from varying *σ* values (***↓, ↑***) and the omission (**o**.) of the Gaussian blur. **Red** text color indicates a high effect, **green** a medium, and **blue** text color a low effect. **C, D**: Datasets C and D. Complete set of effect sizes is shown in Table S3.

| measurements | | Glass's $\Delta$ | | |
| --- | --- | --- | --- | --- |
| | | ↓ | ↑ | **o.** |
| skeletons | C | 1.30 | 0.22 | 3.75 |
|  | D | 0.53 | 0.13 | 1.99 |
| segments | C | 1.46 | 0.25 | 3.35 |
|  | D | 0.90 | 0.22 | 1.58 |

Our results show that certain parameters strongly affect the network measurements (Table S3) and, therefore, require special attention during parameter adjustment. Parameters with a strong effect are, for example, the *σ* value for the Gaussian blur and the brightness settings, as well as the Branch Pruning threshold for experiments where the endpoint number is relevant. In contrast, variations in other parameters, such as the Analyze Particles threshold and the maximum and minimum filters, result in less deviation from the optimized data.

The network measurement data generated with default settings are mainly consistent with those produced with the optimized settings, except for the volume fraction measurement (Figure 10, Table 4). This suggests that the default settings can reasonably be used for preliminary screening. However, the process should always be controlled by visual assessment of the returned binary and skeletonized images compared to the raw image.

**Figure 10.**
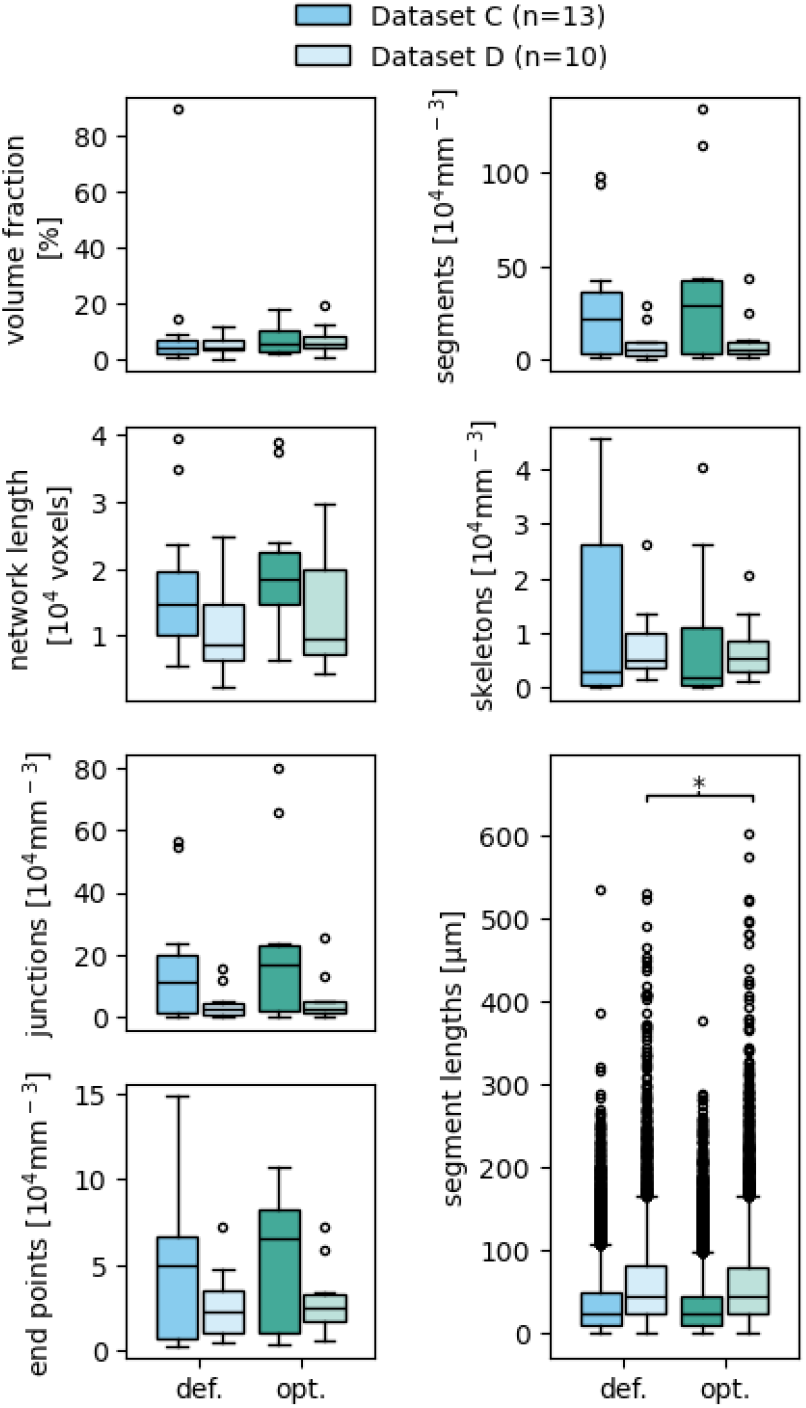
Comparison between the measurement data resulting from default (**def**.) and optimized (**opt**.) parameter settings. Boxes that are marked with an asterisk (*) show a statistically significant deviation for a significance level of *α* = 0.05 between the default and optimized parameter settings, based on a two-sided Mann-Whitney U test. Complete data are shown in Figure S1.

**Table 4.** Effect sizes of the deviation of the measurement data resulting from the default parameter settings from those resulting from optimized parameter settings. **Red** text color indicates a high effect, **green** a medium, and **blue** text color a low effect. **C, D**: Datasets C and D. Complete set of effect sizes is shown in Table S3.

| measurements | | Glass's $\Delta$ |
| --- | --- | --- |
| volume fraction | C | 0.94 |
|  | D | 0.39 |
| network length | C | 0.28 |
|  | D | 0.29 |
| skeletons | C | 0.31 |
|  | D | 0.13 |
| segments | C | 0.22 |
|  | D | 0.17 |
| junctions | C | 0.22 |
|  | D | 0.18 |
| end points | C | 0.19 |
|  | D | 0.19 |
| segment lengths | C | 0.07 |
|  | D | 0.03 |

### 3.3 Application to Sample Data

#### Effect of Anti-Angiogenic Drug on Vasculature

To demonstrate VESNA’s possible application, it was used to assess the effect of sorafenib on the vascular networks generated in blood vessel organoids.

Sorafenib is a tyrosine kinase inhibitor. One of its many functions is the suppression of the activation of the vascular endothelial growth factor receptors VEGFR-1 and -2 that are necessary for vascular development, vessel integrity, and cell contacts [32]. In tumor tissues, the VEGF/VEGFR system is oftentimes deregulated to induce neoangiogenesis, which allows for increased nutrient supply to the tumor cells and was shown to be associated with increased growth and dissemination of the tumor [32, 33]. Due to the antiangiogenic effect of sorafenib, it has been approved for treatment of some carcinomas by the FDA in 2005 [34], and by the EMA in 2006 [35]. We used VESNA to investigate the network sizes in sorafenib-treated organoids

The acquired images of the drug-treated and control groups (Section 2.2, Dataset B) are processed using our macro. Due to the large dimensions, the raw images are scaled by a factor of 0.5 in the x and y dimensions prior to processing. This greatly reduces the necessary processing power and time. The scaling is not expected to lead to a relevant loss of information in the case of this dataset, as the generated vascular networks show sufficiently low detail. Additionally, in one image, part of a second organoid was included in the imaged volume and was manually removed from the image stack. The parameters were iteratively adjusted by visual evaluation of the resulting binary and skeletonized images. The final parameter settings can be found in Table S1. Two of the available images of drug-treated organoids were not able to be properly processed even after parameter adjustment due to high background fluorescence and low fluorescence intensity of the vascular tissue. These images were omitted from further data analysis.

Organoids of the control group and the drug-treated group are depicted in Figure 11a and 11b, respectively. The control group organoid shows a vascular network that is distributed throughout most of the organoid. In contrast, the drug-treated organoid lacks vasculature in a large central area. This observation already aligns with the antiangiogenic effect of sorafenib treatment. The size difference between the two organoids can likely be attributed to random variation, given the considerable heterogeneity of organoid sizes throughout the experiment.

**Figure 11.**
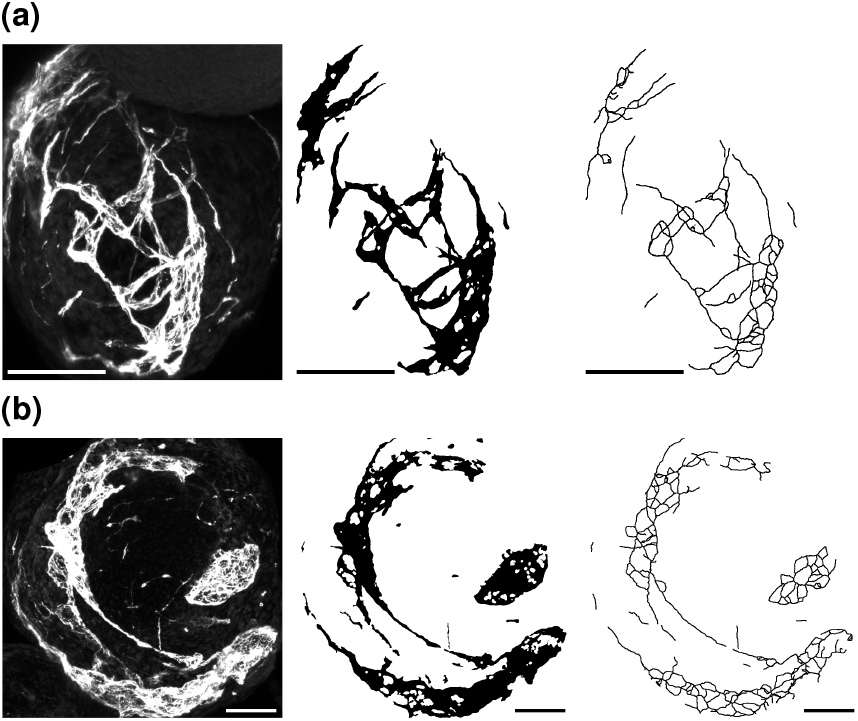
Exemplary cropped Z maximum projections of blood vessel organoids of the control group (**a**) and the group treated with 15 nM sorafenib (**b**) of Dataset B. Depicted are (from left to right) raw images, binary images, and skeletonized images. Skeletonized images are dilated for better visibility. Scale 100 µm.

The resulting network measurement data (Figure 12) show a decrease in all measurements, except for the number of skeletons. While the measurements do not show a statistically significant deviation, in part due to the low numbers of images per group (*n* = 6 for the control group, *n* = 3 for the drug-treated group), the associated Glass’s Δ (Table 5) suggest that sorafenib has a considerable effect on the structure of the vascular networks. The networks seem to be smaller and less complex under drug treatment conditions, as indicated by decreased numbers of junctions and endpoints and shorter networks. Notably, while the segment length diverges in a statistically significant way, the effect size is small in comparison to the other measurements. This can be explained by the high number of data points generated for this measurement, which favors a statistical significance.

**Figure 12.**
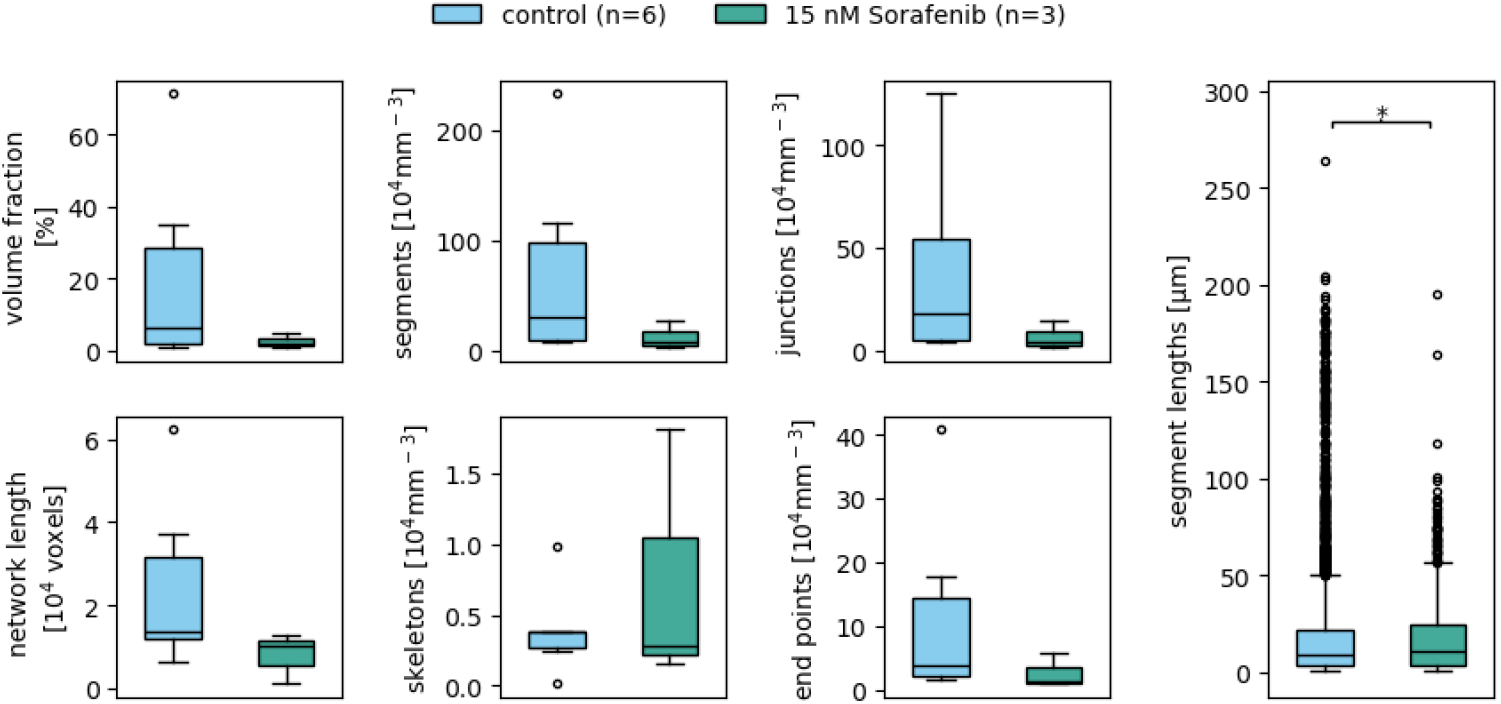
Network measurement data resulting from the drug-treated and control groups of Dataset B. The utilized parameter values are listed in Table S1. Boxes that are marked with an asterisk (**\***) show a statistically significant deviation for a significance level of *α* = 0.05.

**Table 5.** Effect sizes of the deviation of the different network measurement data resulting from the drug-treated blood vessel organoids compared to the control group of Dataset B. **Red** text color indicates a high effect, **green** a medium, and **blue** text color a low effect.

| measurements | Glass's $\Delta$ |
| --- | --- |
| volume fraction | 0.69 |
| network length | 0.82 |
| skeletons | 1.20 |
| segments | 0.72 |
| junctions | 0.73 |
| end points | 0.63 |
| segment lengths | 0.04 |

These observations coincide with the exemplary images (Figure 11). Considering that the measurements are normalized to the imaged volume, the images of the drug-treated organoids show fewer skeletons, junctions, and endpoints than the control group organoid.

The presented results are consistent with the above-mentioned biochemical functions of sorafenib as an antiangiogenic agent and demonstrate the possible application of our macro VESNA to biomedically relevant experimental setups.

#### Effect of Growth Conditions on Vasculature

Another possible application of our macro VESNA is the analysis of vascular networks in organoids generated from varying growth conditions. For this, image data of organoids generated using smaller and larger molds for cell seeding are utilized. The exact growth conditions are detailed in Section 2.2, Dataset C. For the processing of the images, only the brightness parameters are adjusted to accommodate for the low fluorescence intensity of the original images. Parameter settings can be found in Table S1.

Exemplary sections of the images resulting from both groups are depicted in Figure 13. Taking the scales of the images into consideration, the vascular structure of the organoid generated from a small mold (Figure 13a) is more detailed and smaller. On the other hand, the structure generated from seeding in the larger mold (Figure 13b) shows less detail per imaging volume.

**Figure 13.**
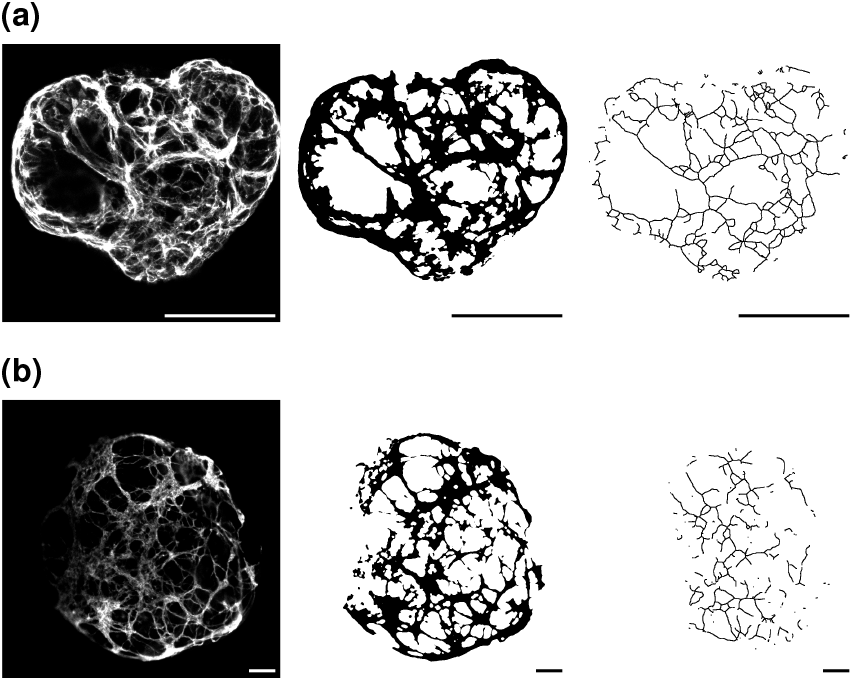
Exemplary cropped Z maximum projections of blood vessel organoids of Dataset C seeded in small molds (**a**) and large molds (**b**). Depicted are (from left to right) raw images, binary images, and skeletonized images. Skeletonized images are dilated for better visibility. Scale 100 µm.

Plotted network measurement data allow a closer look at the difference in structure between both groups of vascular networks. Even at the low number of data used (*n* = 9 for small molds, *n* = 4 for big molds), five out of the seven measurements show statistically significant deviations (Figure 14) and high effect sizes (Table 6). While seeding of the organoids in small molds creates more complex vascular networks with high numbers of segments, junctions, and endpoints, they are also more fragmented, as indicated by the increased number of skeletons. The usage of larger molds leads to simpler networks with fewer but, on average, longer segments. The network length and volume do not differ much. These observations are also reflected in the exemplary sections (Figure 13).

**Figure 14.**
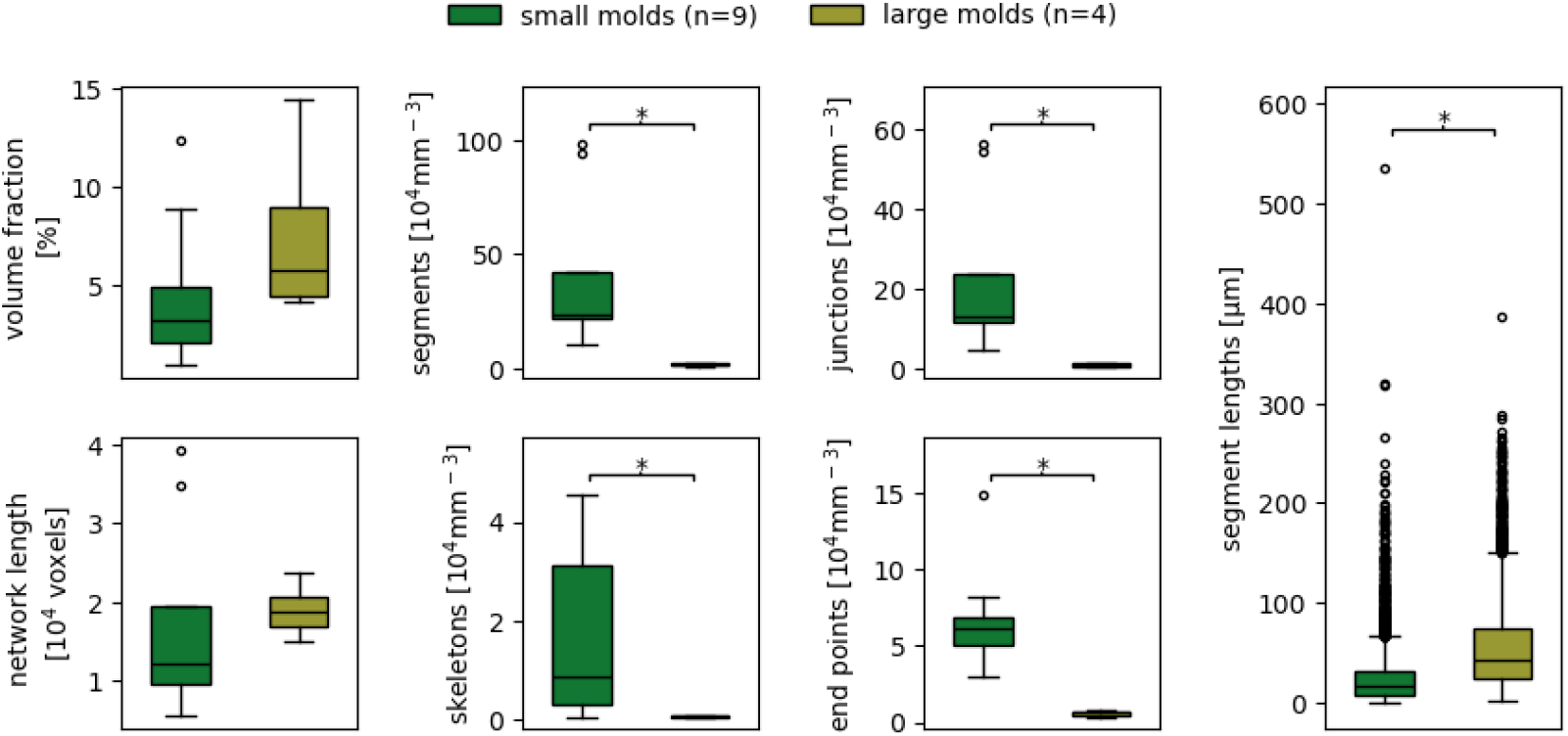
Complete measurement data resulting from the two different mold size groups of Dataset C. The utilized parameter values are documented in Table S1. Boxes that are marked with an asterisk (**\***) show a statistically significant deviation for a significance level of *α* = 0.05.

**Table 6.** Effect sizes of the deviation of the different network measurement data resulting from the different mold sizes of Dataset C. **Red** text color indicates a high effect, **green** a medium, and **blue** text color a low effect.

| measurements | Glass's $\Delta$ |
| --- | --- |
| volume fraction | 0.87 |
| network length | 0.18 |
| skeletons | 1.24 |
| segments | 1.10 |
| junctions | 1.19 |
| end points | 1.84 |
| segment lengths | 1.19 |

#### Application to Hydrogel-Based Cultures

The usage of our macro VESNA is not only limited to microscopy images of organoids, but can easily extend to other three-dimensional culturing methods. To demonstrate this, we processed and analyzed images of three-dimensional hydrogel-based blood vessel cultures (Section 2.2, Dataset D).

Similar to the processing of the images of the organoids described above, the parameters for the processing of the hydrogel culture images were acceptable for the most part, only needing optimization in the brightness parameters (Table S1).

The exemplary image sections show a vascular network at day 7 (Figure 15a) that consists of a majority of shorter segments. The older network at day 17 (Figure 15b) is mainly made up of longer segments, but there are still a number of small segments to be found. Notably, the segments in the structure at day 7 seem to be randomly oriented in all directions, while the vessels at day 17 show a more parallel arrangement (Figure 15).

**Figure 15.**
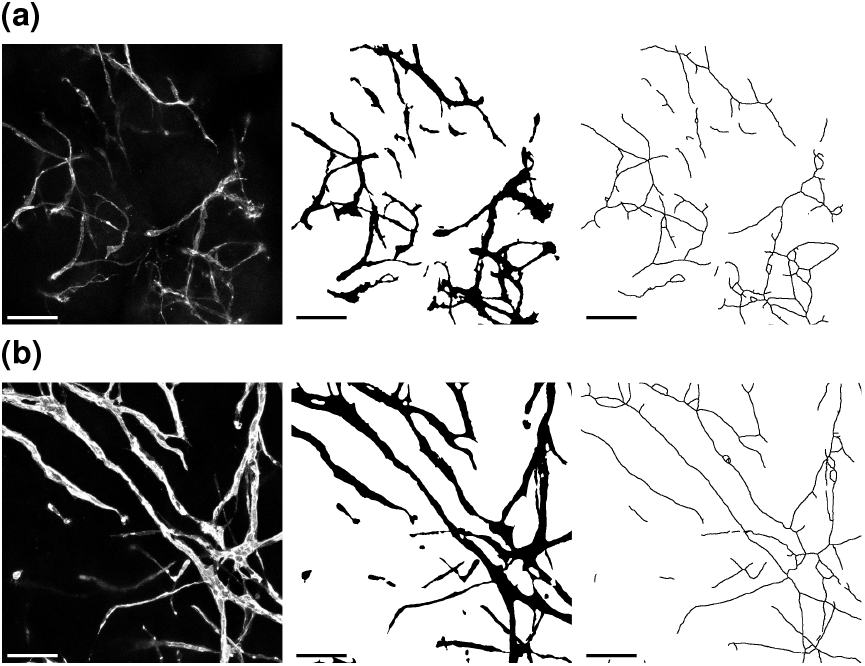
Exemplary cropped Z maximum projections of hydrogel-based blood vessel cultures of Dataset D at days 7 (**a**) and 17 (**b**). Depicted are (from left to right) raw images, binary images, and skeletonized images. Skeletonized images are dilated for better visibility. Scale 100 µm.

The limited sample size at day 7 (*n* = 2) likely accounts for the absence of statistical significance in most measurements and may also contribute to the relatively high effect sizes observed (Figure 16, Table 7). Despite this, the data suggest growth in vascular networks between days 7 and 17, as evidenced by increased segment lengths. While the median segment length remains similar, vascular structures at day 17 exhibit an increased upper limit of segment lengths. This growth is further supported by elevated counts of skeletons, segments, junctions, and endpoints, along with increases in vascular length and volume fraction, all of which are also observed in the exemplary images (Figure 15).

**Figure 16.**
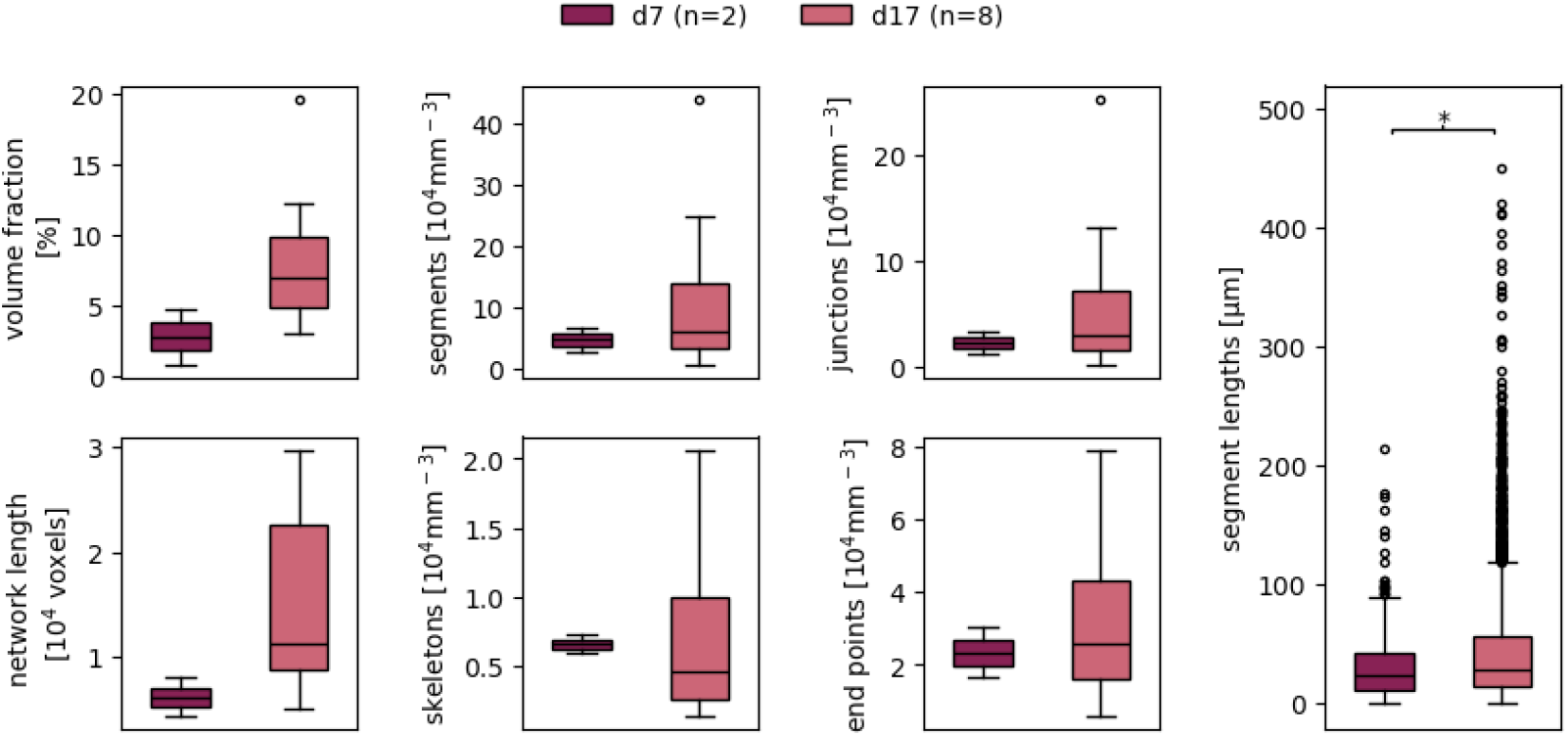
Complete network measurement data resulting from hydrogel cultures of Dataset D at days 7 and 17. The utilized parameter values are documented in Table S1. Boxes that are marked with an asterisk (**\***) show a statistically significant deviation for a significance level of *α* = 0.05.

**Table 7.** Effect sizes of the deviation of the different network measurement data resulting from hydrogel-based cultures of Dataset D at days 7 and 17. **Red** text color indicates a high effect, **green** a medium, and **blue** text color a low effect.

| measurements | Glass's $\Delta$ |
| --- | --- |
| volume fraction | 2.85 |
| network length | 4.85 |
| skeletons | 3.62 |
| segments | 1.11 |
| junctions | 3.40 |
| end points | 1.36 |
| segment lengths | 0.36 |

Our results highlight the versatility of VESNA in facilitating quantitative analyses of vascular networks in organoids and hydrogel-based cultures across diverse research applications, including studies on drug treatments and mechanical constraints.

## 4. Discussion

### 4.1 VESNA’s performance

A visual comparison of the generated skeletonized images to a manually labeled ground truth demonstrates that the segmentation quality is generally very high, with some weaknesses observed in regions exhibiting low signal intensity. Quantitative analysis is particularly sensitive in this context, given the thinness of the skeletonized structures. Despite this sensitivity, the overlap between the skeletonized image and the ground truth proves robust across a range of parameter settings. The adjustment of the *σ* value for the Gaussian blur and the brightness settings yield the largest effects, even though they are still small.

Most network measurements remain consistent and reliable under varying parameter values. High effects of parameter adjustment are observed for the *σ* value of the Gaussian blur, the brightness settings, and the Branch Pruning threshold, indicating that these parameters need to be selected carefully. Smaller effects are observed for parameters such as the Analyze Particles threshold and the maximum and minimum filters. Through this analysis, we underlined the robustness of our macro and identified the most critical parameters for optimizing the segmentation and measurement process, providing guidance for applications of VESNA.

The optimized parameter settings yield network measurement data comparable to those obtained with the default settings. This suggests that fully automated processing using the default parameters is feasible for data sets with similar fluorescence intensity, background fluorescence, and levels of detail as our test data. This ability to bypass the time-intensive parameter optimization process further enhances the macro’s applicability and usability across diverse research scenarios.

### 4.2 Applications

The current version of our macro does not include a representation of vessel direction, which limits its ability to distinguish between random and statistically significant observations in this case. This highlights a possible area of further improvement for our macro. (?)

By demonstrating VESNA’s performance across three different applications, we provide evidence for its robustness across different image data and experimental objectives. First, we tested the effect of sorafenib on the vascular structure generated in blood vessel organoids. Among other mechanisms of action, sorafenib inhibits the activation of VEGFRs, which are responsible for proliferation, angiogenesis and vascular sprouting, explaining the drug’s anti-angiogenic effect [32, 33]. This is consistent with our observed results, showing reduced network size and complexity, as indicated by a decrease in junction and endpoints, as well as number of skeletons compared to the control group. Kumar *et al*. [36] were able to show similar results in a 2D chick chorioallantoic membrane-based experiment. They observed a decrease in network length, branch length, and junction point counts in sorafenib-treated chicken embryos compared to the control group. In opposition to our results, they have also observed a decrease in branch numbers, whereas we observed an increase. We have also tested the effect of different growth conditions on the vascular networks and found that organoids generated from smaller molds develop more complex but also more fragmented vascular structures as evidenced by increased numbers of segments, junction and endpoints, and skeletons. An alternative explanation for the difference in measurements might be the different staining behavior of larger organoids, which is known to be a potential cause of decreased signal intensity towards the center of the organoid [37]. This low fluorescence intensity is also present in our corresponding sample image (Figure 13b), supporting the existence of this effect in our case. But while this effect might seem like a reasonable alternative explanation, the sample image also shows that our macro is able to successfully recognize these low intensity structures. Instead, a likely explanation for the different vascular structures resulting from different growth conditions might be very similar: Analogous to how fluorophore-coupled antibodies are limited in their diffusion to the center of a large organoid, leading to lower signal intensity, VEGF is likely also diffusion-limited to some degree. This would result in an effectively lower concentration of VEGF in the center of larger organoids compared to smaller organoids, restricting the level of activation of VEGFRs and the resulting angiogenesis.

Finally, we demonstrate the ability of our macro to process images generated from other 3D tissue culturing methods, with an example of hydrogel-based cultures. In this experiment, we observe an increase in all measurements as the vascular network grows from day 7 to 17, as expected. The application to other tissue cultures is enabled by the simplicity of our macro’s requirements towards the input images. Specifically, it only requires a vasculature-specific stain, such as CD31, and in the case of larger objects, tissue clearing is also necessary. This provides flexibility that not only makes the processing of a wide range of blood vessel cultures possible, but, given an appropriate antibody target, makes the processing of lymphatic networks or even other vascular-like networks plausible.

Collectively, these three example applications demonstrate our macro to exhibit robust usability across different experimental goals, as well as across a diverse range of image data generated from different tissue culturing methods and varying in features such as age, network size and complexity, fluorescence intensity, background noise, as well as image size. All three datasets are successfully processed after minor parameter adjustment, indicating an even wider range of possible image data than shown here.

In summary, VESNA offers automated image processing for vascular network structures with a wide range of applications. Its straightforward and accessible source code allows for easy customization to meet unique research needs, leveraging the extensive library of macros and plugins available for Fiji. For example, a wider range of image data might be processed by including or adjusting pre- and postprocessing steps. The inclusion of further measurements such as vessel direction would allow for even deeper insight into vascular development. VESNA represents a valuable tool for researchers, streamlining image analysis and opening up new possibilities in the study of vascular development, disease mechanisms, and drug discovery.

## 5. Acknowledgments

SE, PW and LD acknowledge the financial support provided by the TRR 225 Biofab (DFG), Project B04. ChatGPT and Grammarly have been used to improve wording and spelling.

## Appendix

**Table S1.** Default parameter values preset in the macro and chosen values for processing of each set of sample data. Dashes (**-**) indicate no deviation from default values.

| parameters | default | Dataset A | Dataset B | Dataset C | Dataset D |
| --- | --- | --- | --- | --- | --- |
| brightness min | 0 | - | - | 1 | 1 |
| brightness max | 120 | 110 | 255 | 50 | 50 |
| Gaussian blur<br>$\sigma$ [voxels] | 1.8 | - | 2.0 | - | - |
| size threshold<br>[pixels] | 55 | - | - | - | - |
| maximum filter<br>[pixels] | 3 | - | - | - | - |
| minimum filter<br>[pixels] | 4 | - | - | - | - |
| pruning length<br>[voxels] | 20 | - | - | - | - |

### Additional Data on Sensitivity of Parameters

**Table S2.** Parameter values used for the processing of the images of Datasets A, C and D in the parameter sensitivity test. The selected parameter values are chosen to lie just outside of the value range that was found to be useful for most datasets. Dashes (**-**) indicate no deviation from optimized values, while **o**. indicates the omission of the function.

| parameters | def. | opt. | Brightness |  |  |  | Gaussian blur |  |  | Analyze Particles |  |  | Max. and Min. filters |  |  | Branch Pruning |  |  |
| --- | --- | --- | --- | --- | --- | --- | --- | --- | --- | --- | --- | --- | --- | --- | --- | --- | --- | --- |
|  |  |  | min ↑ | max ↓ | max ↑ | o. | ↓ | ↑ | o. | ↓ | ↑ | o. | ↓ | ↑ | o. | ↓ | ↑ | o. |
| brightness min | 0 | 1 | 30 | - | - | o. | - | - | - | - | - | - | - | - | - | - | - | - |
| brightness max | 120 | 50* | - | 30 | 200 | o. | - | - | - | - | - | - | - | - | - | - | - | - |
| Gaussian blur<br>$\sigma$ [voxels] | 1.8 | 1.8 | - | - | - | - | 0.5 | 3.0 | o. | - | - | - | - | - | - | - | - | - |
| size threshold<br>[pixels] | 55 | 55 | - | - | - | - | - | - | - | 20 | 150 | o. | - | - | - | - | - | - |
| maximum filter<br>[pixels] | 3 | 3 | - | - | - | - | - | - | - | - | - | - | 1 | 6 | o. | - | - | - |
| minimum filter<br>[pixels] | 4 | 4 | - | - | - | - | - | - | - | - | - | - | 1 | 7 | o. | - | - | - |
| pruning length<br>[voxels] | 20 | 20 | - | - | - | - | - | - | - | - | - | - | - | - | - | 10 | 100 | o. |

**Figure S1.**
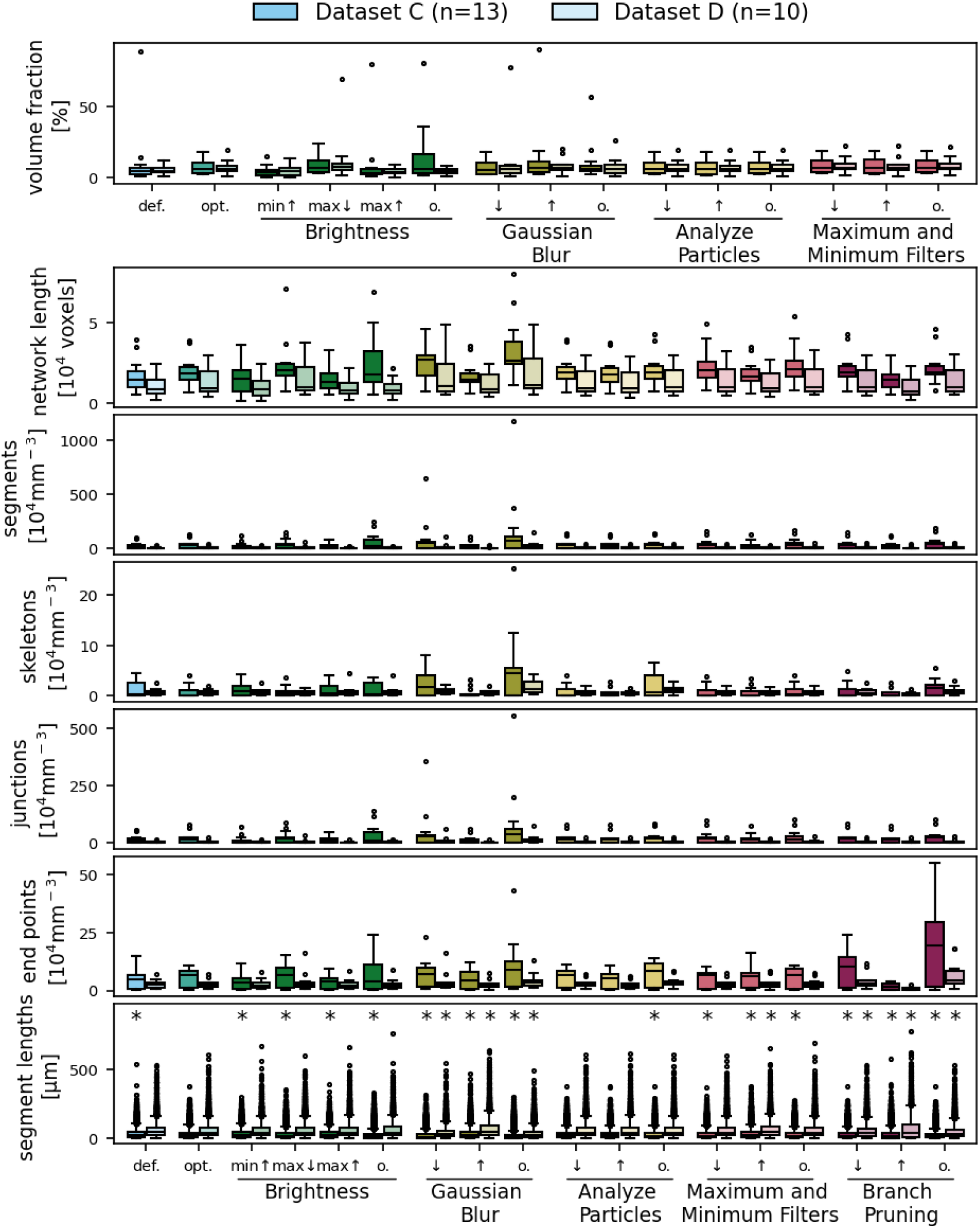
Complete measurement data resulting from the parameter sensitivity test performed on images of Datasets C and D. The utilized parameter values are documented in Table S2. Boxes that are marked with an asterisk (*) show a statistically significant deviation from the optimized measurement data, using a Bonferroni-corrected significance level *α* = 0.0036 for the volume fraction measurement and *α* = 0.0029 for the remaining measurements.

**Table S3.**
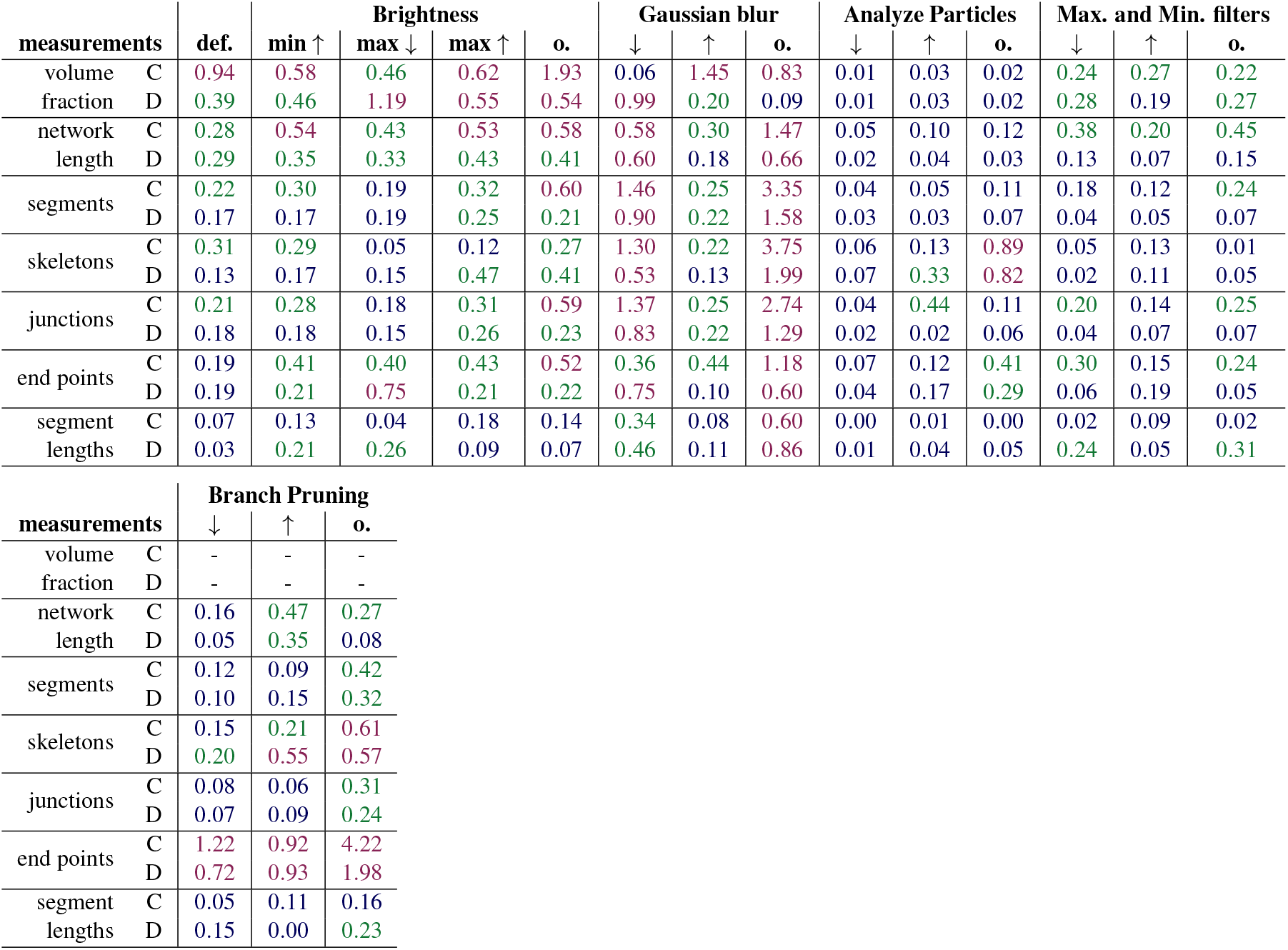
Effect sizes (rounded to two decimal places) of the deviation of the resulting measurement data of tested parameter settings from the optimized settings. **Red** text color indicates a high effect, **green** a medium, and **blue** text color a low effect. **C, D**: Datasets C and D.

